# Revisiting animal photo-identification using deep metric learning and network analysis

**DOI:** 10.1101/2020.03.25.007377

**Authors:** Vincent Miele, Gaspard Dussert, Bruno Spataro, Simon Chamaillé-Jammes, Dominique Allainé, Christophe Bonenfant

**Affiliations:** Université de Lyon, F-69000 Lyon; Université Lyon 1; CNRS, UMR5558, Laboratoire de Biométrie et Biologie Évolutive, F-69622 Villeurbanne, France; CEFE, Univ Montpellier, CNRS, EPHE, IRD, Univ Paul Valéry Montpellier 3, Montpellier, France; Mammal Research Institute, Department of Zoology & Entomology, University of Pretoria, Pretoria, South Africa; LTSER France, Zone Atelier “Hwange”, Hwange National Park, Bag 62, Dete, Zimbabwe

**Keywords:** individual identification, deep metric learning, image similarity networks, open-source software

## Abstract

An increasing number of ecological monitoring programs rely on photographic capture-recapture of individuals to study distribution, demography and abundance of species. Photo-identification of individuals can sometimes be done using idiosyncratic coat or skin patterns, instead of using tags or loggers. However, when performed manually, the task of going through photographs is tedious and rapidly becomes too time consuming as the number of pictures grows.

Computer vision techniques are an appealing and unavoidable help to tackle this apparently simple task in the big-data era. In this context, we propose to revisit animal re-identification using image similarity networks and metric learning with convolutional neural networks (CNNs), taking the giraffe as a working example.

We first developed an end-to-end pipeline to retrieve a comprehensive set of re-identified giraffes from about 4, 000 raw photographs. To do so, we combined CNN-based object detection, SIFT pattern matching, and image similarity networks. We then quantified the performance of deep metric learning to retrieve the identity of known individuals and detect unknown individuals never seen in the previous years of monitoring.

After a data augmentation procedure, the re-identification performance of the CNN reached a Top-1 accuracy of about 90%, despite the very small number of images per individual in the training data set. While the complete pipeline succeeded in re-identifying known individuals, it slightly under-performed with unknown individuals.

Fully based on open-source software packages, our work paves the way for further attempts to build automatic pipelines for re-identification of individual animals, not only in giraffes but also in other species.

## 1 Introduction

In many respects, population and behavioural ecology have immensely benefited from individual-based, long term monitoring of animals in wild populations (Clutton-Brock & Sheldon, 2010; Hayes & Schradin, 2017). At the heart of such monitoring is the ability to recognize individuals. This is often achieved by actively marking animals, such as deploying ear-tags or leg rings, cutting fingers or feathers, or scratching scales in reptiles (Silvy *et al*., 2005). In some species, however, individuals display natural marks that make them uniquely identifiable. For instance, many large African mammals such as leopard (*Panthera pardus*), zebra (*Equus sp*.), kudu (*Tragelaphus strepsiceros*), wildebeest (*Connochaetes taurinus*) or giraffe (*Giraffa camelopardalis*), all present idiosyncratic fur and coat patterns particularly useful for non-invasive and reliable recognition of individuals. Individual identification in the wild has long been known to be feasible from comparisons of the distinctive coat patterns of individuals (Estes, 1991). As the number of individuals to identify increases, people-based visual comparisons of pictures can rapidly become overwhelming. With the recent move to digital technologies (namely digital cameras and camera traps), the problem becomes even more acute as the number of pictures to process can easily reach the thousands or ten of thousands.

Over the last decade, the use of computer vision rapidly spread into biological sciences to become a standard tool in animal ecology for many repetitive tasks (Weinstein, 2018). In a seminal publication, Bolger *et al*. (2012) first presented computer-aided photo-identification, initially for giraffes but more recently applied for dolphins (Renó *et al*., 2019). The underlying computer technique is a feature matching algorithm, the Scale Invariant Feature Transform operator (SIFT; Lowe (2004)), where each image is associated to the *k*-nearest best matches. The current use of SIFT for ecologists requires human intervention to validate the proposed candidate images within a graphical interface (Bolger *et al*., 2011). In the same vein, other feature-based proposals were developed in the last decade to apply computer vision on different types of idiosyncrasies (Hartog & Reijns, 2014; Moya *et al*., 2015). A drawback of the method frequently arises when two images are considered similar not because of similar skin or coat patterns of animals, but because of similarities in the backgrounds (presence of distinctive tree for instance), hence leading to false positive matches. For the best results with computer vision, all images should be cropped before, so that only the relevant part of the animal appears in the images (for instance, excluding most of the neck, head, legs and background for large herbivores). Until now, this cropping operation was most often done manually (Halloran *et al*., 2015), despite being a highly time-consuming task when processing thousands of images.

Meanwhile, the Deep Learning (DL) revolution was underway in computer vision, showing breakthrough performance improvements (Christin *et al*., 2019). In particular, convolutional neural networks (CNNs) are now the front-line computer technique to deal with a large range of image processing questions in ecology and environmental sciences (Lamba *et al*., 2019). Many recent studies tackle the general problem of re-identification using CNNs, which has been mostly developed and extensively used for humans (Wu *et al*., 2019). Technically, re-identification consists in using a CNN to classify images of different individuals, some of them being not necessarily seen before, *i*.*e*. unknown individuals. However, despite the availability of proven and efficient techniques (Zheng *et al*., 2016), and several successful attempts to apply the method to non-human species (Körschens *et al*., 2018; Hansen *et al*., 2018; Moskvyak *et al*., 2019; Bouma *et al*., 2019; Schofield *et al*., 2019; He *et al*., 2019; Bogucki *et al*., 2019; Schneider *et al*., 2020; Chen *et al*., 2020; Ferreira *et al*., 2020), re-identification remains a challenging task when applied to animals in the wild where re-observations are limited in number to train the model satisfactorily *sensu largo* (Schneider *et al*., 2019).

In practice, current CNN-based approaches have to be tailored to the needs of field ecologists interested in using them for individual recognition. For instance, batches of new images are regularly added to the reference database following yearly fieldwork sessions because of the recruitment of newborns or of immigrants if the study population is demographically open. Therefore, we expect the re-sighting of known individuals, as well as the observation of individuals never seen before. In other words, this standard sampling design implies to solve the re-identification in a mixture of known and unknown individuals. Chen *et al*. (2020) referred to this problem as the “open set” identification problem, and they proposed to identify images from unknown individuals and to assign them a single “unknown” label. In practice, those unknown individuals are important to identify because they are new animals entering the database from which the life history will be built on.

A classical CNN classifier can re-identify already known individuals (usually with a *softmax* last layer) but will fail to identify new individuals. Indeed, the number of predicted classes must match the number of known individuals. Therefore, we crucially need a CNN-based approach that can filter out individuals unknown at the time of the analysis. We propose to rely on deep metric learning (DML, see Hoffer & Ailon, 2015) as an ideal candidate to solve the “open set” identification problem: DML consists in training a CNN model to embed the input data (input images) into a multidimensional Euclidean space such that data from a common class (for instance, images of a given individual) are much closer, in terms of Euclidean distance, than with the rest of the data. Retrieving the known and unknown individuals consists in relying on the Euclidean distance computed for any pair of images.

Here we addressed the problem of photo-identification with an updated, open-source, and end-to-end automatic pipeline applied to the case for the iconic and endangered giraffe (*Giraffa camelopardalis*). In a first step, we applied state-of-the art techniques for object detection with CNNs (Lin *et al*., 2017) to automatically crop giraffe flanks of about 4,000 raw photographs shot in the field. Indeed, the most recent CNN approaches clearly outperformed other approaches (Girshick *et al*., 2014), including the Histogram of Oriented Gradients (HOG) approach that was recently used with giraffes (Buehler *et al*., 2019). Second, following Bolger *et al*. (2012), we used the SIFT operator to calculate a numeric distance between any pair of giraffe flanks. From the *n × n* calculated distances, we followed the new framework of image similarity network (Wang *et al*., 2018) and applied unsupervised learning to retrieve different clusters of images coming from different individuals, hence removing any human intervention in the process. However, we manually validated a subset of our results to build a ground-truth data set of different individuals (*n* = 82). Using this data set as a training set, we finally developed a supervised learning strategy using CNNs and evaluated its predictive accuracy with a cross-validation procedure.

## 2 Material and Methods

### 2.1 Photograph database

We carried out this study in the northeast of Hwange National Park (HNP), Zimbabwe. HNP park covers a 14,650 km^2^ area (Chamaillé-Jammes *et al*., 2009). The giraffe sub-species currently present in HNP could be either *G. c. angolensis* or *G. c. giraffa* according to the IUCN (Muller *et al*., 2018). Here we used data from a regular monitoring of individuals conducted between 2014 and 2018. Each year for at least three consecutive weeks, we drove the road network daily within *<*60km of the HNP Main Camp station, and took photographs of every giraffe encountered. Pictures were taken with 200mm to 300mm lenses mounted on Nikon DSRL cameras (sensor resolution ranged between 16 and 40 Mpx). At this stage we filtered sequences of very similar photographs occurring with the camera burst mode in the same second, and retained one single photograph per sequence. Overall, we retained *n* = 3,940 photographs.

### 2.2 Image cropping with CNN and transfer learning using RetinaNet

A range of CNN-based tools are now available for object detection, already used for animal detection (Parham *et al*., 2018; Schneider *et al*., 2018; Sadegh Norouzzadeh *et al*., 2019). Among other options including YOLO (Redmon *et al*., 2016; Bochkovskiy *et al*., 2020) and Mask R-CNN (He *et al*., 2017), RetinaNet (Lin *et al*., 2017) is a CNN-based object detector able to detect a series of predefined object classes (*e*.*g*. different animal species) that returns the coordinates of a bounding box around these objects, and a confidence score as well. These two steps are performed at the same time with a single CNN, which makes RetinaNet a *one-stage* detector as opposed to two-stage detectors for which a first CNN search for regions containing a potential object and a second CNN classify these regions Redmon *et al*. (2016). However, RetinaNet better manages non informative objects’ background with similar performance compared to two-stage detectors while being much faster (Lin *et al*., 2017). Finally, it is known that the more heterogeneous the training data set is (various positions, backgrounds, scale or lighting), the most efficient a CNN is (Beery *et al*., 2018), so we used data augmentation (flipping, rotation and color changes) to enhance our model performance.

For an efficient detection and classification of objects, a CNN has to be trained on a huge amount of images (usually *>* millions of images) to capture the most discriminant features associated with each class. Because of the limited number of photographs we have at hand, we relied on *transfer learning* (Shin *et al*., 2016). Transfer learning is a specific method aiming at training a CNN on a small number of images that do no start CNN training “from scratch” with some random model parameters, but uses the parameters of a model pre-trained on a large data set and for similar tasks as the one of interest (Willi *et al*., 2019). This approach works because the pre-trained model has already learnt a wide range of relevant and generic features.

We manually prepared our training data set by cropping bounding boxes around giraffe flanks, excluding most of the neck, head, legs and background, with the `labelImg` open source program for image annotation (https://github.com/tzutalin/labelImg). We obtained 469 bounding boxes associated to a subset of 400 photographs. We performed transfer learning with RetinaNet to detect a single object class, the giraffe flank, from a pre-trained model shipped with RetinaNet, that is a ResNet50 backbone trained on the COCO dataset (80 different classes of common objects including giraffes among a few other animal species; see Lin *et al*. (2014)). We trained the model with 30 epochs of 100 batches of size 2. Our pipeline was based on the Keras implementation of RetinaNet available at https://github.com/fizyr/keras-retinanet.

### 2.3 Identification of individuals using unsupervised learning

#### 2.3.1 Using the Scale Invariant Feature Transform operator

We built on Bolger and colleagues Bolger *et al*. (2012) to achieve pattern matching between giraffe flanks with the Scale Invariant Feature Transform operator (SIFT; Lowe (2004)), currently the most commonly used computer vision approach to identify individuals (Bellavia & Colombo, 2020). The SIFT algorithm extracts characteristic features in photographs called *key points* that are invariant with respect to scale and orientation. Comparing two photographs, pairs of matching key points (*i*.*e*. having similar characteristics) are retrieved and ranked by distance (Euclidean distance between their feature vectors). Here, we selected the 25 closest pairs of key points. However, for better results, we had to assess the extent to which matching key points were coherent in the two giraffe flanks, *i*.*e*. if their location matched on the giraffe body. To find out relevant cases where matching key points were actual matches of coat patterns, we superimposed key points extracted from a pair of giraffe photographs with a geometrical transformation called *homography*. An homography is a perspective transformation between two planes, one for each image, that finds key points from the first image as close as possible from those of the second image. The homography preserves the relative positioning of key points but changes the perspective, i.e. the distance between points. Once the key points were retrieved from the 2 images, they were superimposed on a plane to compute the Euclidean distance between all pairs of key points in a pair of photographs, hence obtaining our SIFT-based distance. We used the implementation of SIFT and homography in the open source `openCV` library version 3.4 (Bradski, 2000).

### 2.4 Image similarity network, community detection and clusters of images

Following the computation of distances between all pairs of giraffe flanks obtained with the SIFT operator approach, we searched for clusters of flank images that should come from one single individual giraffe. We first defined a network made of nodes and representing giraffe flank images, and of edges: we considered that two nodes were connected by an edge, *i*.*e*. two flanks were similar and came from the same giraffe individual, if the SIFT-based distance between paired images felt below a given threshold (see below for more details). Therefore, the so-called *connected components* of this network should associate images from different individuals.

We estimated the distance threshold value by taking advantage of a property of complex networks called the *explosive percolation* (Achlioptas *et al*., 2009). The explosive percolation predicts a phase transition of the network just above a threshold point. At this point, adding a small number of edges in the network, for example by slightly increasing the distance threshold (Hayasaka, 2016), leads to the sudden appearance of a *giant component* encompassing the majority of nodes. In other words, at some point a small increase of the distance threshold leads to considering that almost all images come from the same individual giraffe. We determined the threshold value graphically, selecting the transition point where the giant component starts to increase dramatically (Supp. Fig. fig:giant).

An additional issue arose when different nodes were erroneously connected. This was the case when the distance computation failed in avoiding false positive edges (example in Figure S1), *i*.*e*. when two flanks are erroneously considered similar. Moreover, in some cases the body of two or more giraffes could overlap in one photograph. In this situation, two or more nodes might be linked by edges, when we actually had different giraffes. Therefore, we applied a network clustering algorithm called *community detection*, developed in network science (Fortunato, 2010), to split – only when relevant – any connected component into different groups of nodes that are significantly much more connected between themselves than with the others, a so-called *community*. Indeed, the presence of many edges inside a group of images suggested it was consistent and taken from the same individual, whereas the absence of many edges between two groups clearly informed about their inconsistency and heterogeneity (*i*.*e*. from two different individuals). We applied the community detection with the InfoMap algorithm (Rosvall & Bergstrom, 2008). The final product of the community detection algorithm was a set of *clusters* of images corresponding either to a connected component or to a community retrieved by InfoMap.

### 2.5 Re-identification of individuals, using supervised learning

#### 2.5.1 Deep metric learning and triplet loss with CNN

The principle of deep metric learning is to find an optimal way to project images into an Euclidean space such that the Euclidean distance can be used for machine learning tasks. In this context, we trained a CNN model using the triplet loss (Hermans *et al*., 2017), in line with recent studies on other species Moskvyak *et al*. (2019); Bouma *et al*. (2019). The triplet loss principle relies on triplets of images composed by a first image called *anchor* and another *positive* image of the same class (same giraffe here) and a third *negative* image of another class (any different giraffe) (see Bouma *et al*., 2019, for details). The training step consists in optimizing the CNN model such that the Euclidean distance computed using the last CNN layer (hereafter called CNN-based distance) between any anchor and its positive image is minimal, while maximizing the distance between this anchor image with its negative counterpart. We used an improved algorithm called *semi-hard triplet loss* (Schroff *et al*., 2015), that deals only with triplets where the positive and negative images are close (in other words, the “hard” cases), using the TripletSemiHardLoss function in TensorFlow Addons. After training completion, we computed the Euclidean distances between any pair of giraffe flank photographs, again using the vector composing the last layer of our CNN model.

#### 2.5.2 Data augmentation, training and test data sets

The training and test data sets required for the CNN approach were derived from the photograph clusters identified by the SIFT algorithm. We retained only those clusters fulfilling the following conditions: (i) the cluster contains a minimum of two sequences of images shot at least 1 hour apart; (ii) the cluster can be divided into a first set of sequences large enough to perform training (we imposed at least five images), and a second set of sequences; (iii) the cluster demonstrated a perfect and verified consistency. We used the first set of sequences for CNN training, and the second as an independent test data set to assess the model performance. The first condition ensured that we have complete independence between training and test data sets, i.e. giraffes being seen under different conditions (time, season or location).The third condition is of upmost importance because errors in the data set would lead to sub-optimal performances of the machine learning approach. We therefore carefully checked, manually, that the SIFT-based clusters we used in the CNN were perfectly unambiguous. We achieved this high level of data quality by discarding all cases where two or more giraffes overlapped on the same frame, or when giraffes were indifferently oriented from the back to the front (orientation ambiguities).

We cropped all flank images to focus on the central part of the flank, keeping 80% of the original width and 60% of the height (in particular excluding the neck and its background). By doing so, we wanted to prevent our CNN model from capturing background noise. Additionaly, we homogenized contrast of images by normalizing the three colour channels using the `Imagemagick` package (`normalize` option; https://imagemagick.org). In the end, all images were resized to 224×224 pixels.

We ended up with five flanks per individual at least, and a median of seven (Table 1) in the training set. This particularly low number of images available to train the CNN led us to consider the few shot learning framework, a class of problems where only a few images are available for training. We implemented a 10-fold data augmentation procedure where we made extensive use of image augmentation using the `imgaug Python` library (https://github.com/aleju/imgaug). For each image in the training data set, we performed a random set of transformations such as modifying orientation and size, adding blur, performing edge detection, adding gaussian noise and modifying colors or brightness (details in the available Python code). We finally used this set of eleven images per original image, i.e. the original one and ten modified versions of this image.

**Table 1.** Average number of images and sequences (i.e. separated by at least one hour interval) per individual in train, test and unknown data sets. Median and range, over 10 trials.

|  | Nb. images | Nb. indiv. | Nb. images<br>per indiv. | Nb. sequences<br>per indiv. |
| --- | --- | --- | --- | --- |
| Train | 503 [479-529] | 62 | 7 [5-24] | 2 [1-5] |
| Test | 121 [118-126] | 62 | 2 [1-5] | 1 [1-4] |
| Unknown indiv. | 40 | 20 | 2 | 2 |

### 2.6 Evaluation of CNN-based re-identification

To quantify the overall predictive performance of our CNN deep metric learning, we replicated the following procedure ten times. We first randomly drawn 25% of our individuals from our data set and we considered these are *unknown* individuals. Then for each of them, we randomly selected two images, one in each of the two sequences. This data set is therefore made of individuals seen twice with at least one hour interval between observations, meaning we had two photographs per unknown individuals to work with. With this data set, we aimed to test the ability of the CNN model to detect unknown individuals. The remaining 75% individuals were equivalent to the known giraffes in our population, that is individuals already seen in a previous field session. For these known individuals, we selected all photographs from the first sequence (see above) and used it to built a training data set for the CNN. We kept all images from the remaining sequences as the test data set for *known* individuals. We ensured here a very good independence between training and test data, mostly thanks to the one hour (at least) time lag between observations.

Once the selection of individuals was completed, we performed transfer learning using the pre-trained model ResNetV2 readily available in Keras. We estimated the model parameters using the augmented training data set with 80 epochs with batches of size 42. We used the stochastic gradient descent optimizer with a rate of 0.2. Our pipeline was implemented with Keras 2.3.0.

To mimic re-identification *per se*, literally re-seeing known individuals, we considered that we had a “reference book” with five *representative* images per known individuals: these images were randomly drawn out of the training data set. We then calculated the CNN-based distance between these representative images and each image from the test data set. In essence, we expected small distances between test images and representative ones when they came from the same known individual. Similarly, we calculated the CNN-based distance between representative images and images of the so-called unknown individuals. We also considered that two images can come from the same individual if their distance was below a given threshold. This threshold was a stringency condition that we made vary between 0 and 1.

We quantified the predictive performance of the trained CNN model for different distance thresholds. First, we computed *Top-*1 accuracy for known individuals, consisting in checking for each query image if a representative image from the same individual was the one with smallest distance (*i*.*e*. the Top-1 image) and with a distance below the threshold. In the following, *Top-*1 accuracy was also called *true positive* (TP) rate. Then, we computed the *false positive* rate (FP), checking cases where the Top-1 image was from a different individual. Finally, we quantified the CNN ability to sort out images from unknown individuals. For different distance threshold values, we checked if unknown individual images had a Top-1 image below the threshold. If not, we considered that we successfully detected an unknown individual, hence computing the *true negative* (TN) rate.

## 3 Results

### 3.1 From thousands of photographs to thousands of images of giraffe flank

We trained the object detection method with RetinaNet (Lin *et al*., 2017) on a set of 400 photographs for which the cropping of the giraffe flank has been previously done manually. Training took approximately 30 minutes on a Titan X card. When applying the automatic cropping procedure on our 3,940 photographs (see Figure 1a), we retrieved 5,019 images with associated bounding boxes, supposed to contain a single giraffe flank (see Figure 2a). The cropping failed for 186 photographs (failure rate: 4.7%), mostly due to foreground vegetation and, unusual and difficult orientation of giraffes in the photograph (see examples on Figure 1b). In a few cases, a bounding box could contain the bodies of two overlapping giraffes, one being partially in front of the other (see Figure 2a). Similarly, in some rare instances giraffes were standing very close to each other on a photograph, a situation where RetinaNet could fail in retrieving the exact boundaries of each giraffe flank (see the worst case that we experienced, from a partially blurry photograph in Figure 2b).

**Fig. 1.**
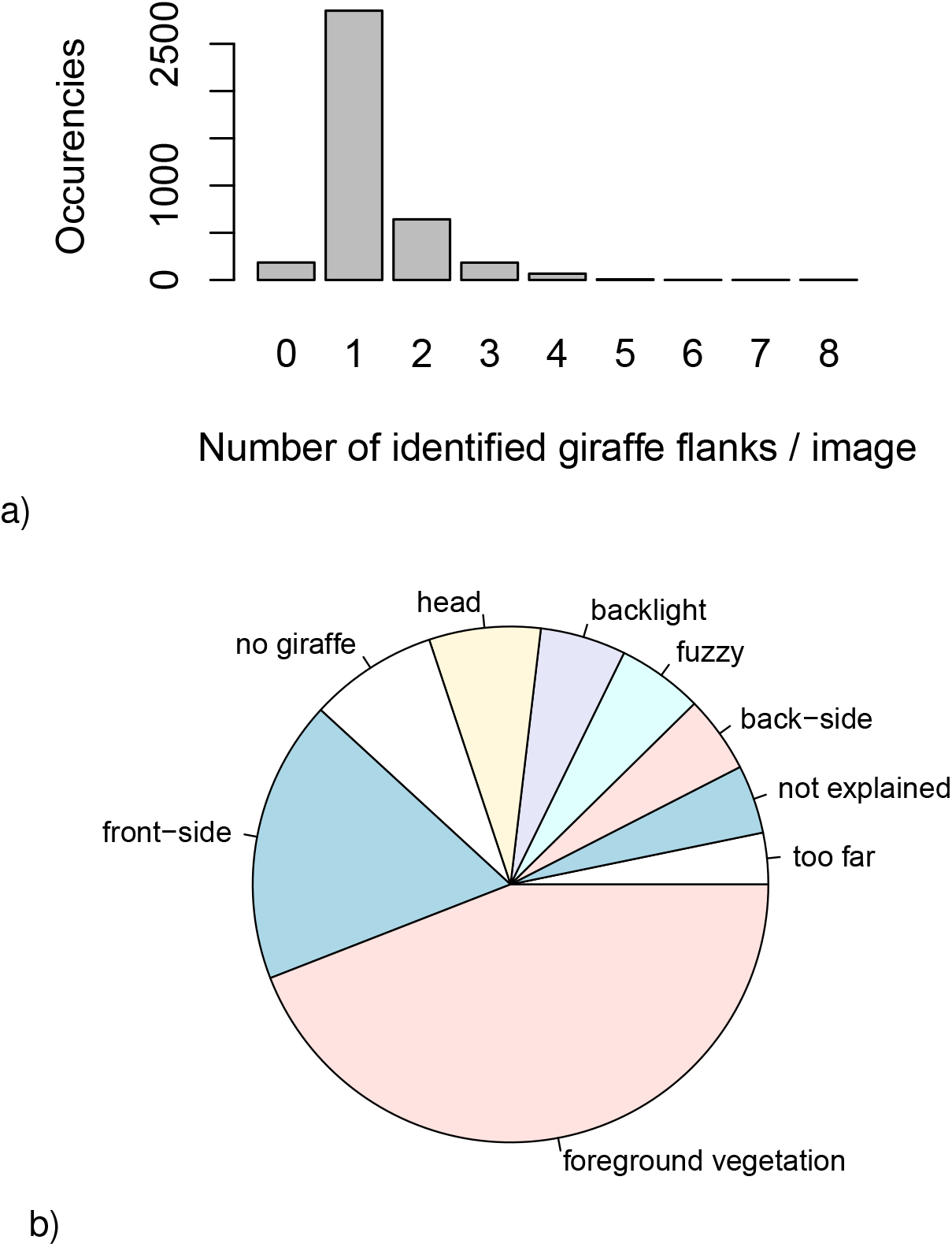
Performance of RetinaNet flank detection. a) Number of identified flanks per image. Manual classification of cropping problems encountered in 186 images where Retinanet failed to identify a giraffe flank.

**Fig. 2.**
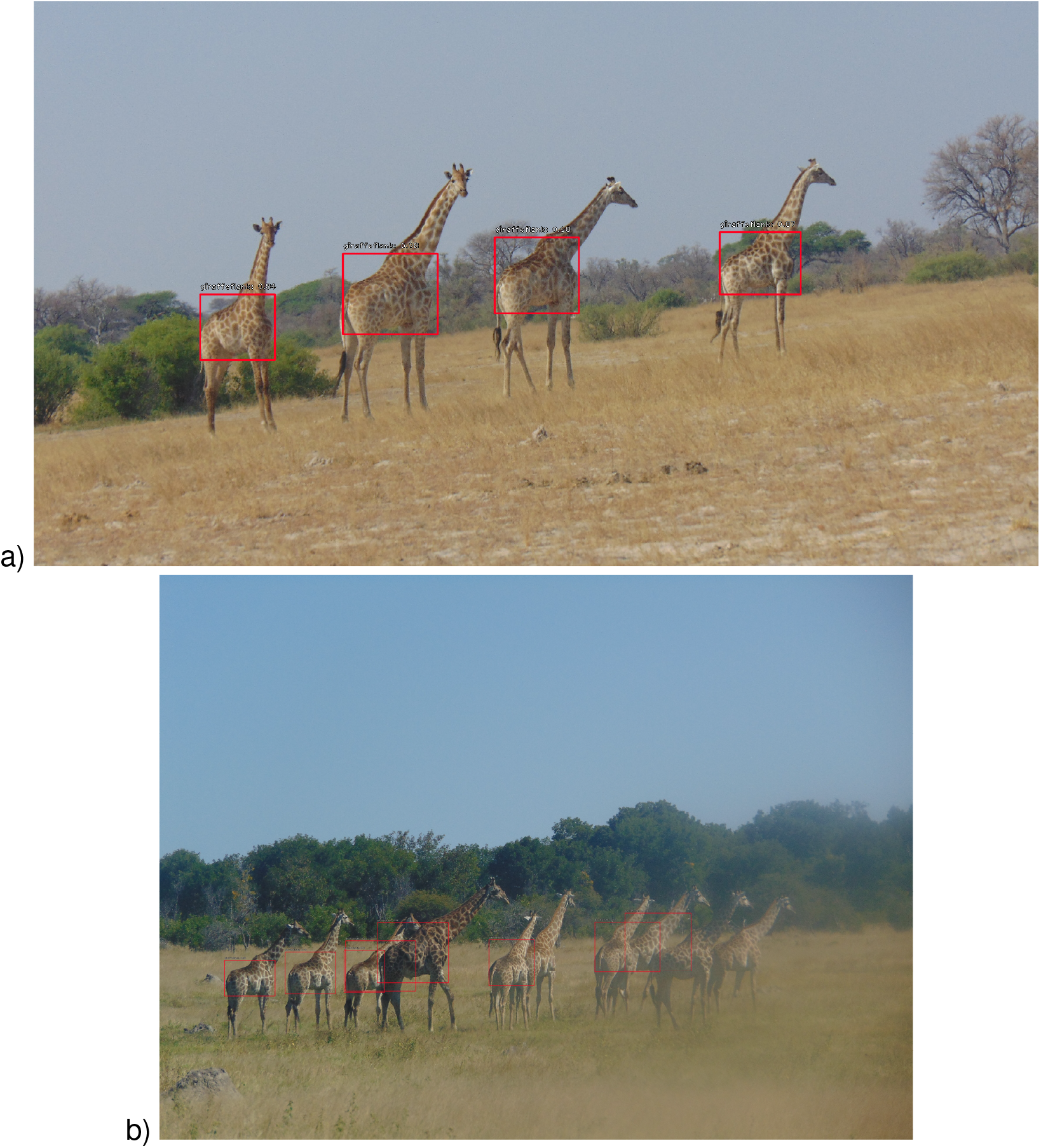
Examples of cropping with RetinaNet. a) Best-case scenario b) Worst-case but rare scenario where different individuals are overlapping and the right-end side of the photo is blurry.

### 3.2 From thousands of images to hundreds of identified individuals

Running the SIFT algorithm (Lowe, 2004) to compare all pairs of flanks took about 800 CPU hours of heterogeneous computing resources. We estimated the threshold value for the giant component (see Methods) at a distance of 340 (see Figure S2a), and obtained an image similarity network composed of 5,019 nodes and 11,249 edges, yielding 1,417 connected components among which 781 were singletons of one image.

Our network-based approach, relying on community detection, retrieved consistent *clusters* of flank images (different colors in Figure 3). The cluster size distribution is by definition more concentrated after network clustering (see Figure S3) with a maximal size of 35 instead of 373. Indeed, this very large connected component was clearly an artifact due to a chain of giraffe overlaps, and has been successfully split by our procedure (see Figure S4). We detected 316 clusters with more than 5 images, and 105 with more than 10 images. However, in rare cases, some images from the same individuals were found in different clusters (see Figure S4). Because these clusters arose from a single connected component, we could *a posteriori* check for consistencies by comparing clusters of the same component manually (such as performed for Figure S4).

### 3.3 From identified individuals to a deep learning approach for re-identification

To perform fair evaluation of the performance, we saved 82 human-validated, unambiguous SIFT-based clusters that contained at least two different sequences of photographs shot at least with a one hour interval (see Material and Methods). Those 82 clusters were made of 822 images of giraffe flanks from which we evaluated the performance of our re-identification pipeline based on deep metric learning. Once trained using data augmentation, the CNN returned a Top-1 accuracy (TP rate) of about 85% on average (Figure 5) for images of known individuals. However, eleven images were found to be repeatedly impossible to classify because of bad orientation of the giraffe body on the photograph, or because of the presence of conspicuous and disturbing elements at the forefront (Supp. Figure S6). Without these problematic images, we achieved a Top-1 accuracy *>*90%, on average. Interestingly, the associated false positive rate was close to 0 (Figure 5). In other words, when a Top-1 image existed below a given threshold (here 1. at most), this Top-1 image was almost always from the correct known individual (Supp. Figure S5 a).

**Fig. 3.**
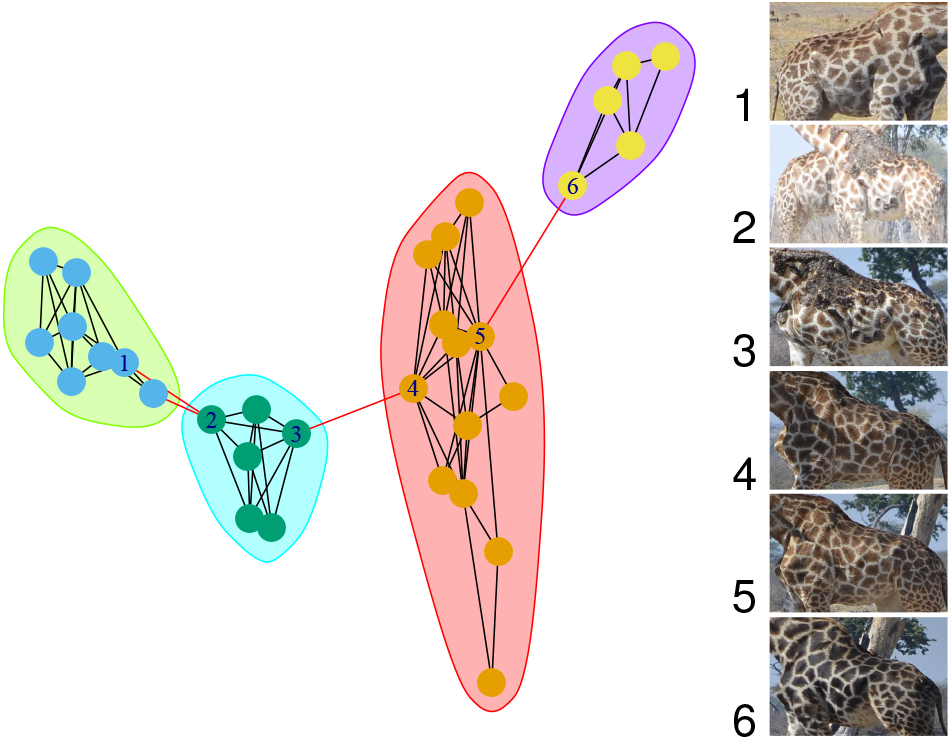
Example of a connected component split into four clusters using the InfoMap algorithm (see Methods). Each cluster is delineated by an ellipse of different color. Node 2 is an image with two giraffes that we also have in images 1 and 3 respectively. Images 3 and 4 are considered similar because of the presence of the same tree in the background (same for images 5 and 6).

**Fig. 4.**
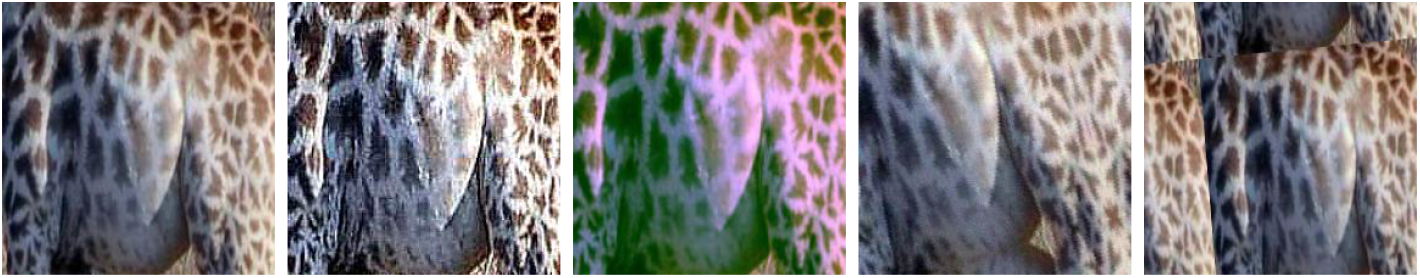
Example of data augmentation with the original image (left) and four different modified versions used to train our CNN.

**Fig. 5.**
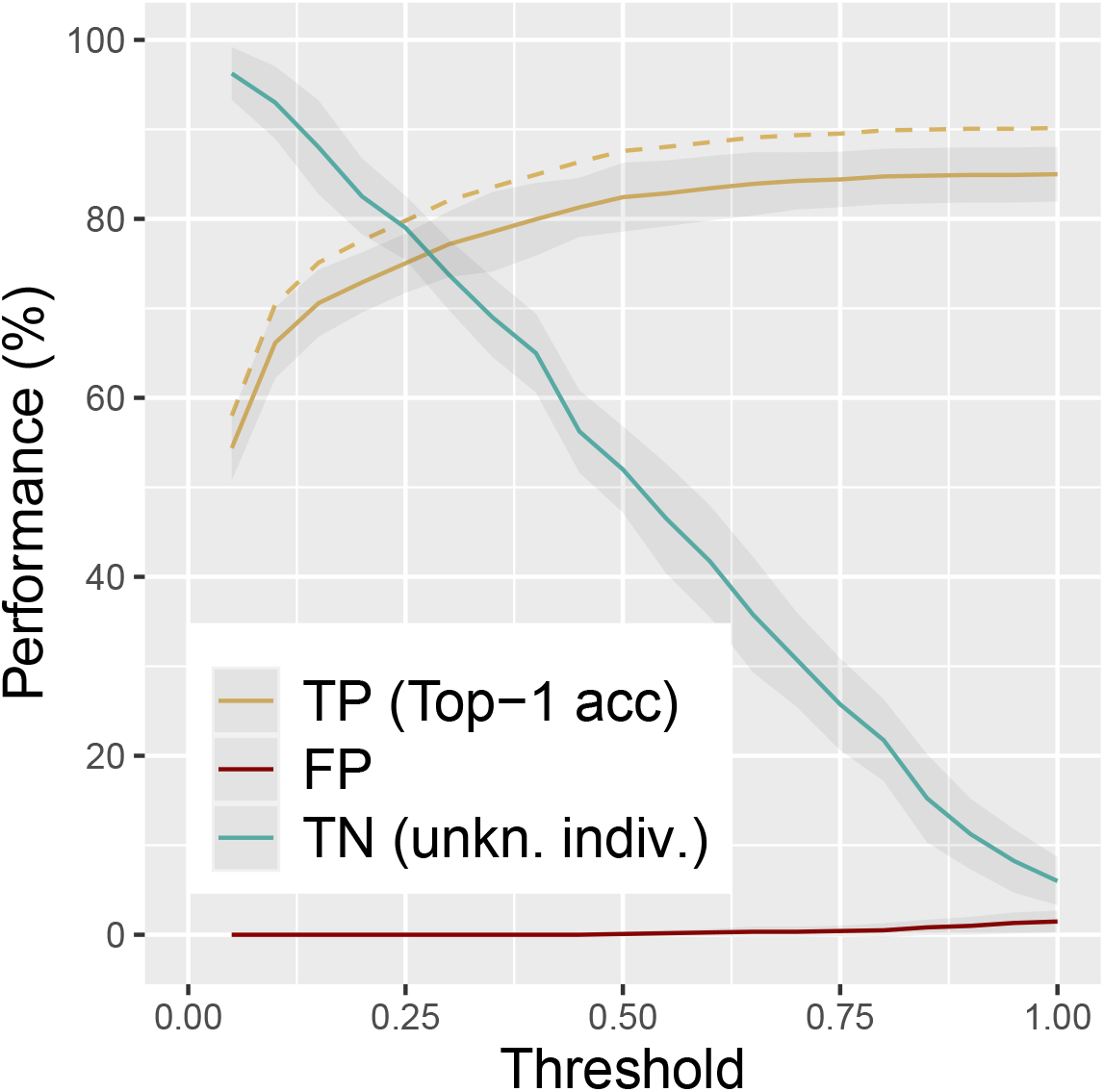
Performance of the re-identification pipeline. True positive rate (TP) or Top-1 accuracy calculated on images of know individuals in the test data set, with (plain) or without (dashed) 11 problematic images. Corresponding false positive rate (FP) or Top-1 error. True negative rate (TP) calculated on images of unknown individuals.

With our deep metric learning approach, images were projected into an Euclidean space. We expected images from the same known individual to be close in this space, whereas images from unknown individuals should be distant from those of known individuals. This prediction was partly supported only. If, for small distance threshold values (*d <*= 0.1) the true negative rate was *>*95%, we found a positive rate *<*70% (Figure 5) and the true negative rate increased with the distance threshold. Hence, our CNN often predicted an unexpected small distance between a given image of unknown individual and another image of a known individual (Supp. Figure S5 b). Interestingly, we observed a particular threshold value (*d* = 0.25) at which both TP and TN rates reached 80% (crossing point in Figure 5), offering the best compromise in terms of correct and false re-identification.

## 4 Discussion

We propose here two complementary approaches to re-identify individual giraffes from a set of photographs taken in the field. Based on the new framework of image similarity networks, our unsupervised method goes one step further compared to previous approaches from the literature since its end product is a comprehensive list of clusters of images, one cluster per identified individuals. Our supervised method, that relies on deep metric learning, achieves a very good re-identification of giraffes from a “reference book” of known individuals despite the rather small number of photographs per individuals available to train the model.

As a first step, we took advantage of the most recent computer vision techniques to perform object detection and crop the giraffe flanks before comparing coat patterns. Image cropping proves to be particularly efficient when the body of several giraffes do not overlap in photographs. However, cascade of problems arises when overlapping occurs, including erroneous cropping and difficulties to assign a bounding box to a single individual because in this case the coat patterns of two individuals are mixed. We show that a limited number of labeled photographs is needed to train RetinaNet (a few hundreds) with a very good performance on new photographs. To what extent our RetinaNet model parameters could be efficient in other study sites with different background vegetation (in “Terra Incognita”, quoting Beery *et al*. (2018)) remains an open question. Nevertheless fine tuning RetinaNet for a particular task and data set is within the reach of many researchers dealing with animal photographs thanks to the associated code we provide. Further perspectives now arise with contour segmentation methods (He *et al*., 2017) than can extract contours of an object such as the whole body or any part of an animal by creating a so-called *segmentation mask* (Brodrick *et al*., 2019). Giraffe body contouring could possibly help for the individual re-identification by removing background residual noise, but building a training set by manually contouring hundreds of animal bodies remains a huge effort.

We then recast the animal identification problem from photographs into a statistical one, namely a clustering problem in an image similarity network. In other words, given a network that we build using a distance between pairs of images, we can efficiently retrieve the image set of a given individual as a cluster in a network. We computed a distance based on pattern matching between flanks with the well known SIFT operator (Bellavia & Colombo, 2020) as used by Bolger *et al*. (2012). The proposed network-based approach was particularly useful and efficient to deal with false positive matches. False positive matches are a recurrent issue occurring when two images have very similar background. This situation is often found when the same tree appears on two images (see nodes 3 and 4 in Figure 3), when giraffe orientation perfectly matches (see Figure S1), or when the bodies of two giraffes overlap on the same image, which is the most frequent configuration we faced (see node 2 in Figure 3). In this latter case, this image linked two sets of images corresponding to the two individuals. Our network-based approach also handles false negative cases (*e*.*g*. differences due to lighting conditions and animal orientation) since community detection is robust to possibly missing edges: indeed, a missing edge can be compensated by the other edges inside a cluster. This step is fully reproducible and applicable to other animal species, as long as a feature matching algorithm can be used, be it SIFT or any other alternative methods such as Oriented FAST and rotated BRIEF (ORB Rublee *et al*., 2011), or deep features (Dusmanu *et al*., 2019; Ma *et al*., 2020)).

We tackled the problem of animal re-identification, literally detecting and identifying previously seen animals, considering that we had a “reference book” with photographs of these known individuals. This fits the needs of field researchers that want to monitor the fate of animals by regularly adding new observations in time, for instance by collecting photographs with camera traps. To do so, we evaluated the possibility to use the rapidly developing convolutional neural networks in a supervised learning framework to achieve deep metric learning. Solving this problem was particularly challenging because of the size of our data set. Previous studies on animal re-identification with CNN indeed relied on a higher number of photographs per individuals (Schneider *et al*., 2020; Ferreira *et al*., 2020). In our case, we had to train the CNN with a few images per individuals only (see Snell *et al*., 2017, on *few shot learning* methods) shot in the field with contrasting environmental and light conditions. This situation corresponds to many field studies, and particularly on large mammals (possibly with the exception of primates), for which population density and animal detection rate are low, limiting the expected number of photograph per individuals. To circumvent this problem, we developed a data augmentation strategy that allowed to increase the variability encountered in the training data set and improved the model performance substantially.

In terms of overall predictive performance, we reached about 90% Top-1 accuracy which is comparable to the previously reported performance in animal re-identification of known individuals (see Schneider *et al*., 2019, for a review) but usually achieved with a much higher number of photographs. The combination of recent deep learning algorithm and data augmentation appears very competitive and efficient, with possible application to difficult practical cases like when working on endangered species. Compared to the more robust SIFT operator, we found that the performance of the CNN is affected by the orientation of giraffe body and noticeably by deviation from perfect side shot. In terms of computing requirements, training our CNN remained time-consuming because the number of images to process is increased dramatically by the data augmentation. This problem is partially counter-balanced by the more computationally efficient calculation of CNN-based distances that increases linearly with the number of photographs (computing one projection per image), compared to the SIFT-based approach for which the computing time is proportional to the square of the number of photographs (computing one matching per image pair). For instance, we got all distances in a minute with the CNN and about two hours with the SIFT operator when applied on the test data set (see Table 2).

**Table 2.** Computing time to compare 310 representative images vers 121 test images (CNN-training with about 5500 images). Intel Xeon CPU E5-2650 v4 2.30GHz (CPU) and Nvidia Titan X card (GPU).

| Task | Avg. computing time |
| --- | --- |
| SIFT-based distance | about 1 hour 45 minutes |
| CNN-based distance | about 1 minute |
| CNN training | about 3 hours 45 minutes (with GPU) |

Our approach was also designed to deal with data sets where known and unknown individuals were present. Dealing with unknown individuals is extremely challenging because no image on these new individuals are available in the training data set. Indeed, most classical CNN-based approaches solve classification problems where the number of classes, the number of individuals for us, was fixed. We showed here that it was possible to filter out unknown from known individuals, while re-identifying a large fraction of known individuals at the same time with a success of 80% (for both TP and TN). However, this trade-off came at the cost of a lower Top-1 accuracy, which we acknowledge is not fully satisfying, as already experienced by other authors (Ferreira *et al*., 2020). Still, in most cases, we could validate the proposed identification by examining the Top-1 for each query image (*i*.*e*. checking its closest image) for both known and unknown individuals. Despite not being fully automated, our CNN approach would require low human intervention.

To what extent the performance of our CNN-based pipeline could be improved with more data? Since it is suitable to any species, further data analysis on other species will help answer this question. However, additional strategies would help including the integration of contextual information (Beery *et al*., 2019; Terry *et al*., 2020) such as time, GPS positioning or animal social context. Using accurate segmentation of animal body (He *et al*., 2017; Brodrick *et al*., 2019) will undoubtedly be a solution against side effects of rectangular cropping. Moreover, this pipeline can be used in an active learning strategy where the machine learning model is assisted by human Norouzzadeh *et al*. (2021). Indeed, using the proposed distance threshold in the Euclidean space, one can iteratively enrich the training data set after manual checking of the most confident Top-1 candidates (below a small distance threshold, to guarantee optimal TN rate) and re-run the estimation procedure.

Finally, this inter-disciplinary work provides guidelines about best practices to collect identification images in the field, if to be used later with an automated pipeline such as the one presented here. Better results can be achieved with simple framing rules of animals with cameras. First the field operator should try to avoid as much as possible overlaying bodies of two or more individuals as this was the most acute issue in our giraffe experience. Note that several but well separated individuals in the same photograph is not a problem at all thanks to the CNN cropping performed as a preliminary stage. Another point to pay attention to is the background which, if too similar on the same images (*e*.*g*. photographs shot from the very same spot) with obvious structures (tree, pond, rocks…) will likely mislead the computer vision algorithm, even on cropped images because cropping is rectangular and do not delineate the animal body. This situation often arises while photographing animals moving in line, as giraffes and many others often do. A last point is the heterogeneity of situations under which animals were observed. We did our best to improve the training data set with data augmentation. However, photographing animals in as many different conditions as possible would most likely improve the results. This includes light conditions (dawn, dusk, noon), orientation of individual or background (open vs. more densely vegetated areas). More specific to CNN re-identification is the need to have a greater number of pictures of photographs per individuals (*>* 50) than what is currently available, so a particular attention should be given, in the field under optimal shooting conditions, to the opportunity to take more photographs of each observed individual.

## Acknowledgments

We thank Jeanne Duhayer for her considerable help in analysing our preliminary findings, and Laurent Jacob and Franck Picard for their insights on deep learning. This work was performed using the computing facilities of the CC LBBE/PRABI. Funding was provided by the French National Center for Scientific Research (CNRS) and the Statistical Ecology Research Group (EcoStat) of the CNRS. We are also grateful to Derek Lee for his kind advice in processing photographs, and for sharing with us his experience in the monitoring of giraffes. Finally, we acknowledge the director of the Zimbabwe Parks and Wildlife Management Authority for authorizing this research, and support from the CNRS Zone Atelier / LTSER program for fieldwork and some of the photographs (collection by P.A. Seeber).

## Authors’contribution

V.M., D.A. and C.B. conceived the study with some inputs from S.C.J. V.M. and G.D. developed the approach and performed the analysis. V.M. and S.C.J. supervised G.D. D.A. and C.B. provided the photographs. B.S. set up the computing architecture. All authors contributed to the manuscript.

## Data Availability

The code to reproduce the analysis is available at https://plmlab.math.cnrs.fr/vmiele/animal-reid/ with explanations and test cases.

## Supporting information

**Fig. S1.**
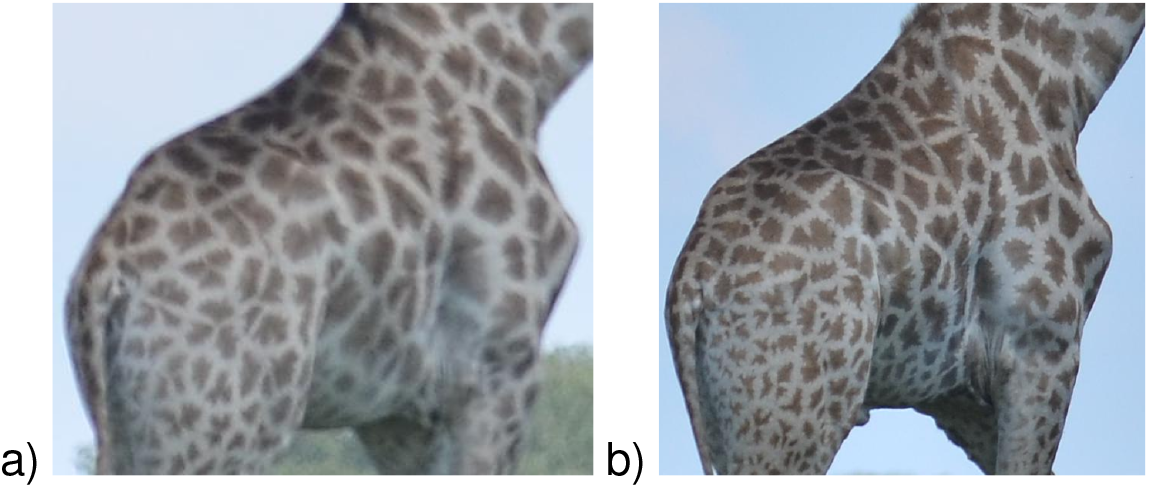
Rare SIFT false positive due to perfect shape and orientation matching. Two different giraffes have a similar pose in a) and b) and the SIFT-based distance between the two images is small and below the used threshold.

**Fig. S2.**
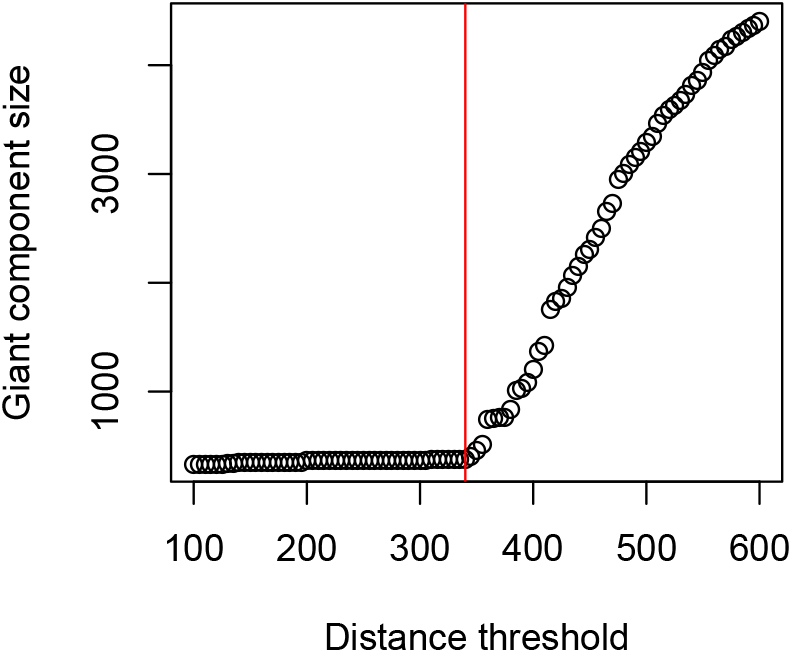
Giant component appearance. We manually estimated the threshold value (red line) used to build our image similarity network. The threshold is 340 when dealing with the SIFT-based distance.

**Fig. S3.**
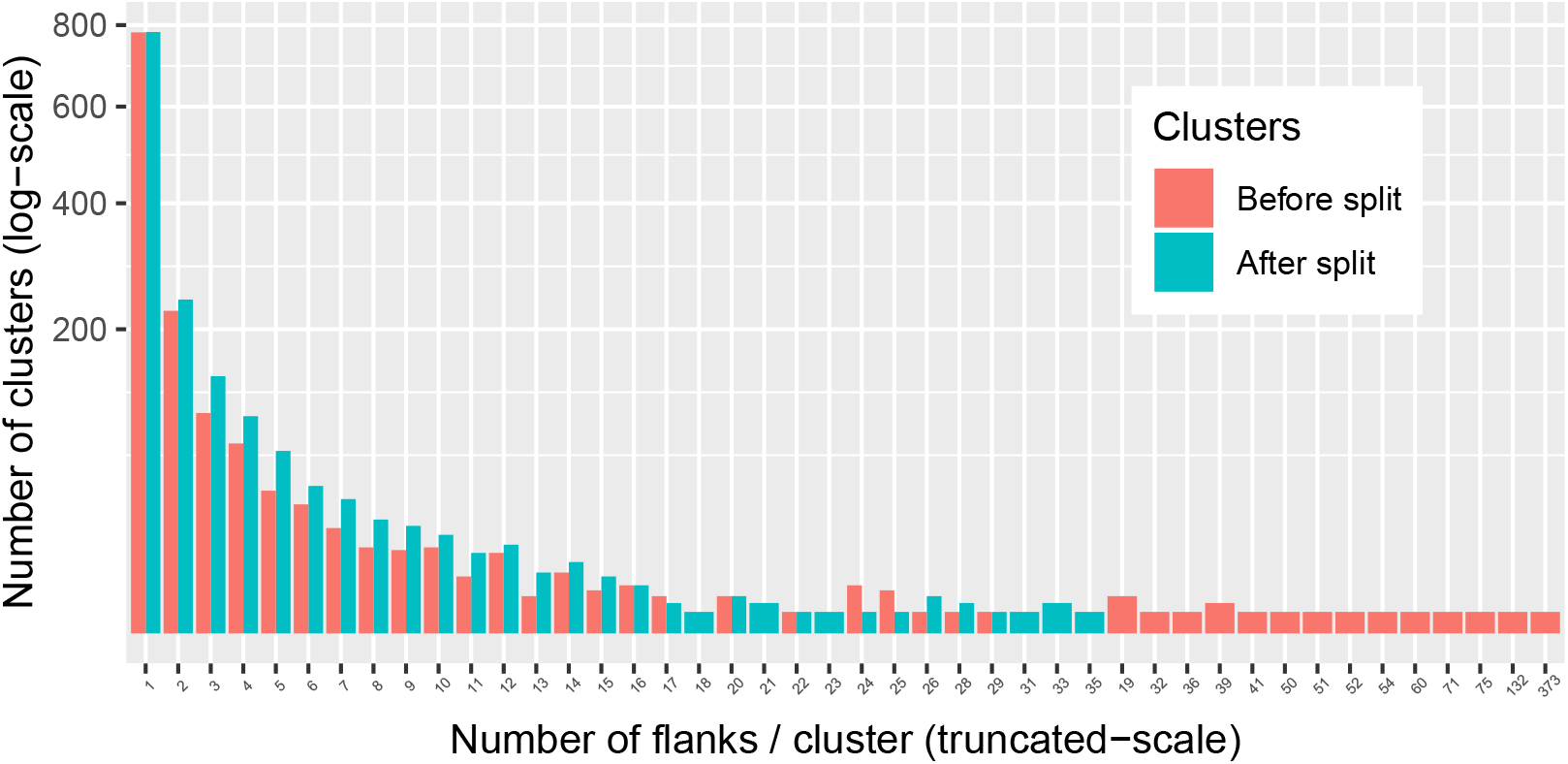
Re-identification from 5,019 giraffe flank images. Number of flank images retrieved by clusters, with the original clusters/connected component (red) or with the clusters retrieved using the InfoMap algorithm to split the connected components (blue; see Methods).

**Fig. S4.**
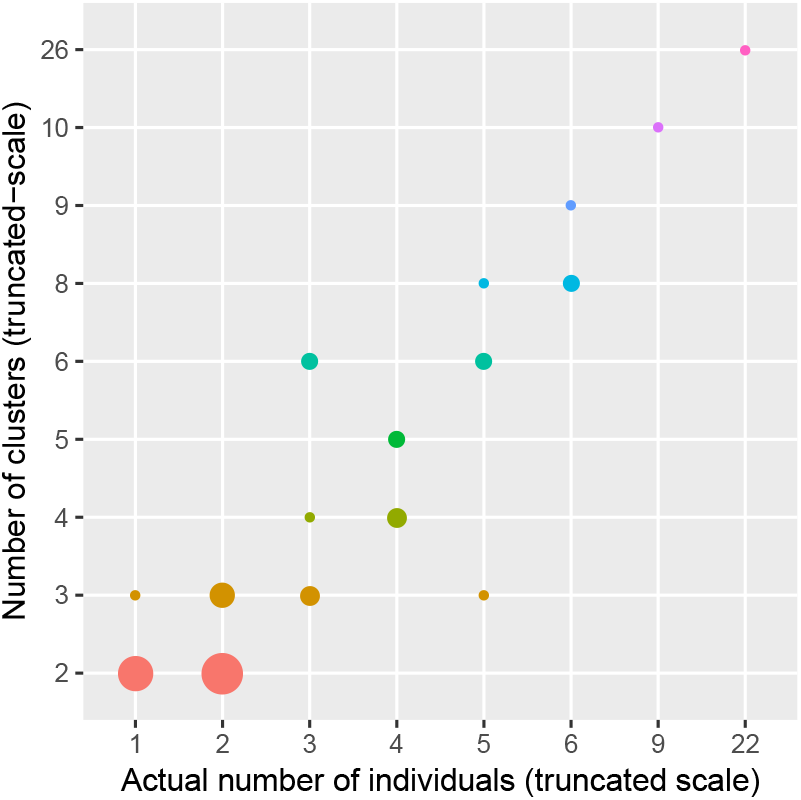
Agreement between the number of clusters (when at least two clusters were found out of a connected component) as returned by our machine-learning approach, and the human-based and manually-checked number of individuals. Circle size is proportional to the number of observations.

**Fig. S5.**
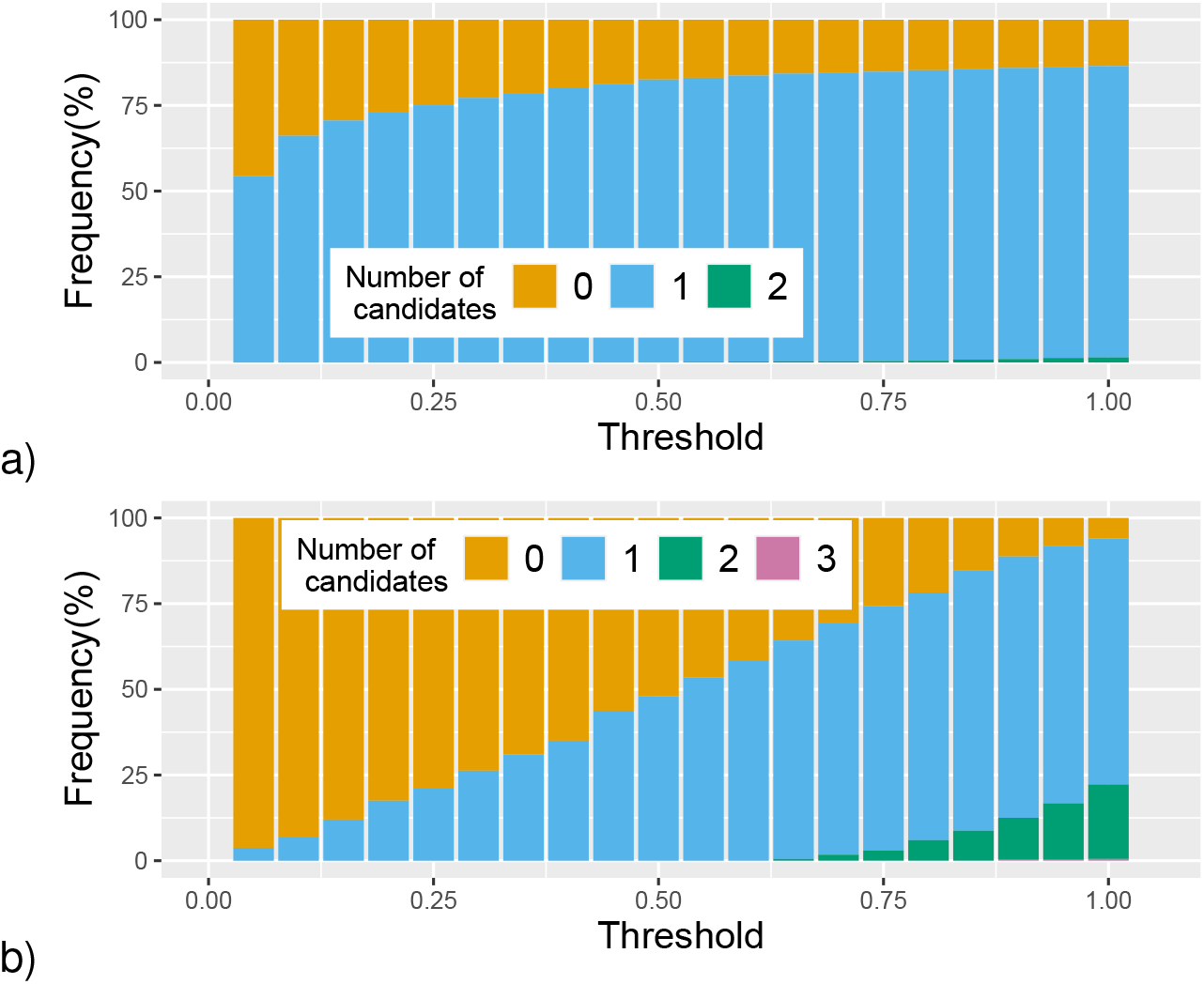
Number of giraffe individual candidates at different distance thresholds. a) Known individuals in the test data set. b) Unknown individuals.

**Fig. S6.**
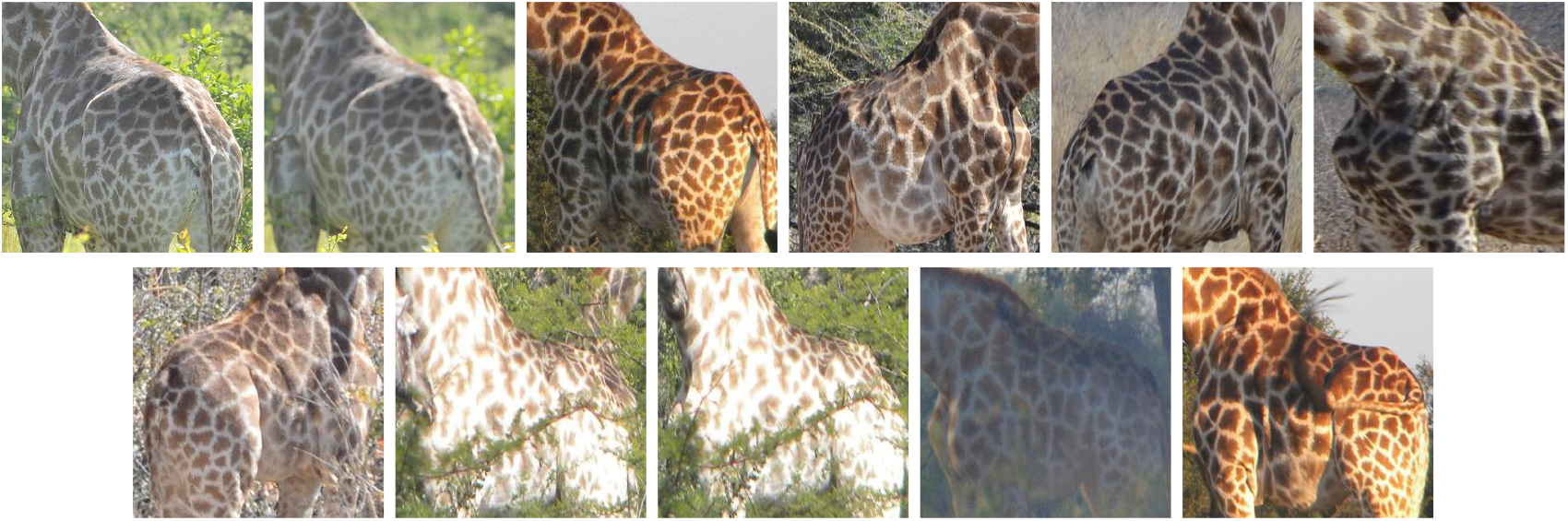
11 problematic images out of the test data set, decreasing Top-1 accuracy because of bad orientation (1st row) or element at the forefront (vegetation or giraffe queue; 2nd row).

## Notes

### Competing Interest Statement

The authors have declared no competing interest.

### Summary of Updates

This version insures a complete independency between train and test data sets.

